# Analysis of Structure and function by the LysR-Type Transcriptional Regulator CbbR of Nostoc sp. PCC 7120

**DOI:** 10.1101/2020.08.04.235895

**Authors:** Hao-Xi Xu, Shu-Jing Han, Yong-Liang Jiang, Cong-Zhao Zhou

## Abstract

The LysR-type Calvin–Benson–Bassham cycle transcriptional regulator CbbR plays an important role in CO_2_ fixation in carbon metabolism in nature, which regulates the gene expression of the key enzyme RibisCO in the Calvin–Benson–Bassham (CBB) cycle. In this study, we optimized the conditions for the transformation, expression, and purification of CbbR in the model algae Nostoc sp. PCC 7120, obtained nick-DNA fragments that could tightly bind to CbbR_7120, and finally obtained CbbR protein crystals. These findings provide great assistance for the final crystallization of CbbR to solve the crystal structure of CbbR, and lay the foundation for understanding the mechanism of CO_2_ fixation in the CBB cycle.

## 1 Introduction

Carbon metabolism is essential for the vital activities of various organisms. Organisms can adapt to different growth environments by delicately regulating the balance of carbon metabolism^[1]^. For most prokaryotes and eukaryotes, CO_2_ is generally the only source of carbon^[2]^. Cyanobacteria are autotrophic photosynthetic microorganisms that have existed on earth since ancient times and are carbon-metabolism model microorganisms to study CO_2_ as the carbon source^[3]^. Most cyanobacteria are aquatic organisms. To adapt to a gradually decreasing concentration of CO_2_, cells have evolved to form a set of CO_2_ concentration mechanisms (CCMs), which can facilitate the acquisition of inorganic carbon source materials (such as CO_2_ and HCO_3_^-^) in cyanobacteria cells under extremely low CO_2_ concentrations^[4]^. The inorganic carbon source is fixed by 1,5-ribulose diphosphate (RuBP) carboxylase/oxygenase (RibisCO) in the carboxysome by the transport system, which increases the CO_2_ concentration in the active center of the enzyme^[5][6]^. In addition, extremely low concentrations of CO_2_ molecules can also be reduced to carbohydrates in the Calvin–Benson–Bassham (CBB) cycle, thereby greatly improving the efficiency of photosynthesis^[7]^.

The ultimate goal of the CBB cycle is to fix three molecules of CO_2_ into one molecule of triose phosphoric acid, that is, the ingested CO_2_ is reduced into the form of available carbon; therefore, these organisms can synthesize the macromolecular structure substances and energy substances essential for vital activities. In cyanobacteria, the uptake of inorganic carbon sources is greatly regulated at the transcription level. CCM-related genes can be induced by low carbon expression, and these genes are inhibited in high-concentration CO_2_ environments^[8]^. The first step of CO_2_ fixation in the CBB cycle is the reaction of CO_2_ with RuBP into 3-phosphoglycerate (3-PGA) under the catalysis of ribulose-1,5-bisphosphate carboxylase oxygenase (RubisCO). The RibisCO-catalyzed reaction is the main pathway for the fixation of inorganic carbon, which is the key enzyme in the C3 pathway and is the center of various enzymes in the CBB cycle. However, due to a lack of high specificity and relatively low catalytic activity, O2 and CO_2_ will compete for the CO_2_ binding site in RibisCO under aerobic conditions; thus, O2 becomes linked to the carbohydrate chain to form the incorrect oxidation products^[9]^. Therefore, cells often increase the total amount of fixed inorganic carbon by synthesizing a large amount of RubisCO^[10]^. Moreover, because the assimilation of CO_2_ has a relatively high energy demand and the burden of additional protein synthesis in cells, the transcriptional regulation of the RubisCo protein gene is of particular significance^[11]^.

The Calvin–Benson–Bassham cycle transcriptional regulator CbbR regulates the gene expression of the key enzyme RibisCO in the CBB cycle^[12]^. The CbbR protein is a type of LysR-type transcriptional regulator (LTTR)^[13]^. CbbR-dependent regulation occurs in different types of organisms, including nonsulfur and purple sulfur bacteria, marine and freshwater chemical autotrophic bacteria, cyanobacteria, methylotrophic bacteria, and different pseudomonas, mycobacteria, fusobacterium, etc. Currently, the spatial structure and function of the CbbR homologous proteins CcmR^[14]^ and CmpR^[15]^ have been partially resolved. These two transcription factors are both members of the LysR transcription factor family^[16]^. This family of proteins can regulate their own recognition of DNA sequences by binding with effector small molecules during the metabolic process^[17]^. However, the relevant structure and specific regulatory functions of CbbR, which is a key regulatory transcription factor that regulates CO_2_ fixation, remain unclarified^[18]^. In the present study, we further analyzed the structure and function of CbbR in future studies by investigating and optimizing the expression, purification, and crystallization conditions of CbbR in model algae.

## 2 Materials and methods

### 2.1 Cloning, expression and purification

The coding region of full-length CbbR was amplified from the genomic DNA of Nostoc sp. PCC 7120 by PCR. The PCR product was cloned into a modified p28 vector with an N-terminal 6× His-tag (Table 1). The recombinant proteins were overexpressed in *Escherichia coli* strain Rosetta cells (DE3). Cells were cultured in LB culture medium (5 g of yeast extract, 10 g of NaCl, 10 g of tryptone per liter, pH 7) containing 16 μg/mL chloromycetin and 30 μg/mL kanamycin at 37°C until the OD_600 nm_ reached 0.8, and then the cells were induced with 0.2 mM isopropyl-β-1-thiogalactopyranoside (IPTG) for another 20 h at 16°C. The cells were harvested by centrifugation at 8 000 g for 4 min at 4°C and resuspended in loading buffer (20 mM Tris, 100 mM NaCl, pH 8.0). After sonication for 15 min and centrifugation at 16 000 g for 30 min at 4°C, the protein supernatant was loaded onto a nickel-nitrilotriacetic acid (Ni-NTA) column (GE Healthcare) containing 0.1 M Ni^2+^ and equilibrated with the same loading buffer. The Ni-NTA column was washed with equilibration buffer. The target protein was eluted with loading buffer containing 400 mM imidazole and then purified by gel filtration with a Superdex 200 column on the ÄKTAprime plus system (GE Healthcare) in binding buffer (20 mM Tris-Cl, pH 8.0, 1 M NaCl, 2 mM DTT, 0.1% β-OG). The purity of the target protein was confirmed by gel electrophoresis. The fractions containing the target protein were pooled and concentrated using a filter tube to a final concentration of 400 mM.

**Table 1.**
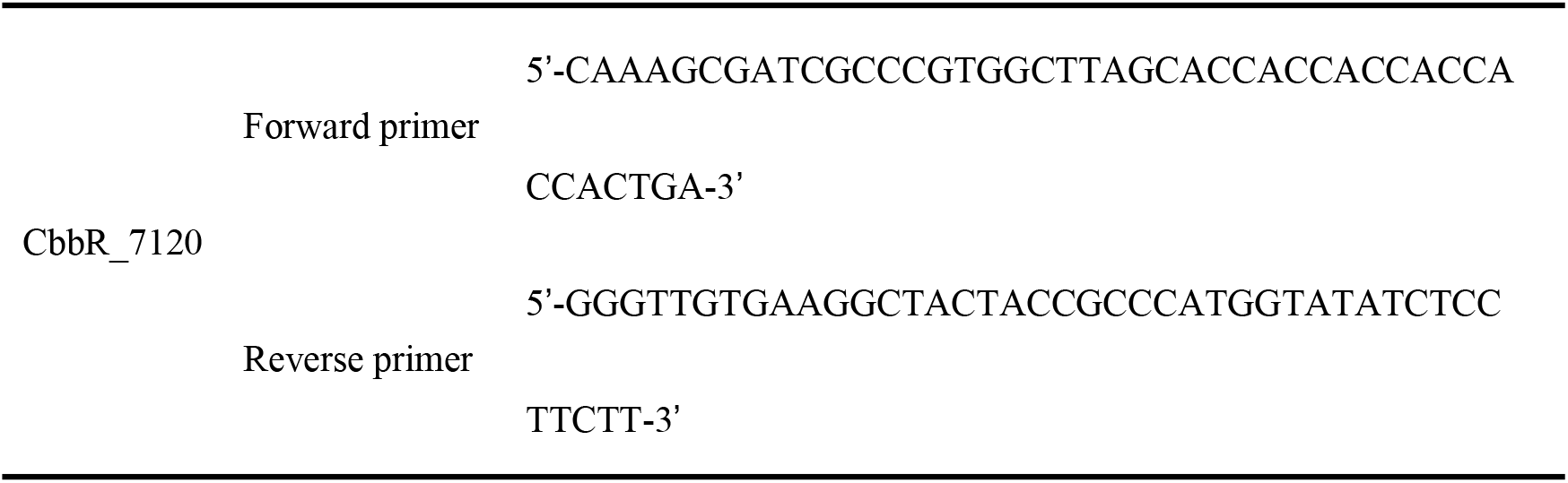
Primer sequence information for the coding region of full-length CbbR from the genomic DNA of Nostoc sp. PCC 7120

### 2.2 Electrophoretic mobility shift assay (EMSA)

EMSAs were performed with 5’-FAM-labeled double-stranded oligonucleotides (dsDNA). The complementary single-stranded oligonucleotides were synthesized by General Biosystems (Chuzhou, China) and annealed to produce double-stranded oligonucleotides. A 10 mL reaction volume containing a 5’-FAM-labeled probe was used with 2.5 μL of 5X binding buffer (100 mM HEPES pH 7.6, 5 mM EDTA, 50 mM (NH_4_)_2_SO_4_, 5 mM DTT, 1% Tween 20, and 150 mM KCl per microliter) and suitable purified recombinant protein. The reaction lasted on ice for 15 min. When required, unlabeled competitors were added to the reaction system for another 15 min on ice. Then, the reaction solution containing protein-DNA complexes was loaded on 6% TBE-polyacrylamide gel with 0.5% TBE loading buffer on ice at 100 V for 2 h.

### 2.3 Crystallization

The full-length CbbR and protein-DNA complexes were applied for crystallization. Crystals were grown at 289 K using a Robot preliminary screen (hanging drop method) on 96-well plates with different conditions.

## 3 Results and discussion

### 3.1 Molecular sieve purification and electrophoresis verification of CbbR_7120

The CbbR protein in Nostoc sp. PCC 7120 model algae was selected. The optimal expression conditions (Table 2) were determined by detecting expression, followed by transformation, massive expression, protein purification and detection according to the optimal conditions.

**Table 2.** The optimal expression conditions of the CbbR protein in Nostoc sp. PCC 7120

| Protein | Competent cells | T (°C) | Position | t (h) |
| --- | --- | --- | --- | --- |
| CbbR_7120 | Rosetta | 16 | Supernatant | 20 |

The protein CbbR_7120 was highly expressed (Fig. 1a). High yield protein bands were present when imidazole was eluted at a concentration of 400 mM but heteroproteins were also present. After purification by molecular sieve chromatography, CbbR_7120 peaked at location D12, and the peak was symmetric and uniform (Fig. 1b). High yield protein bands were present. Peak tip samples were taken for SDS-PAGE to verify protein purity, and we obtained high-purity protein samples (Fig. 1c).

**Figure 1.**
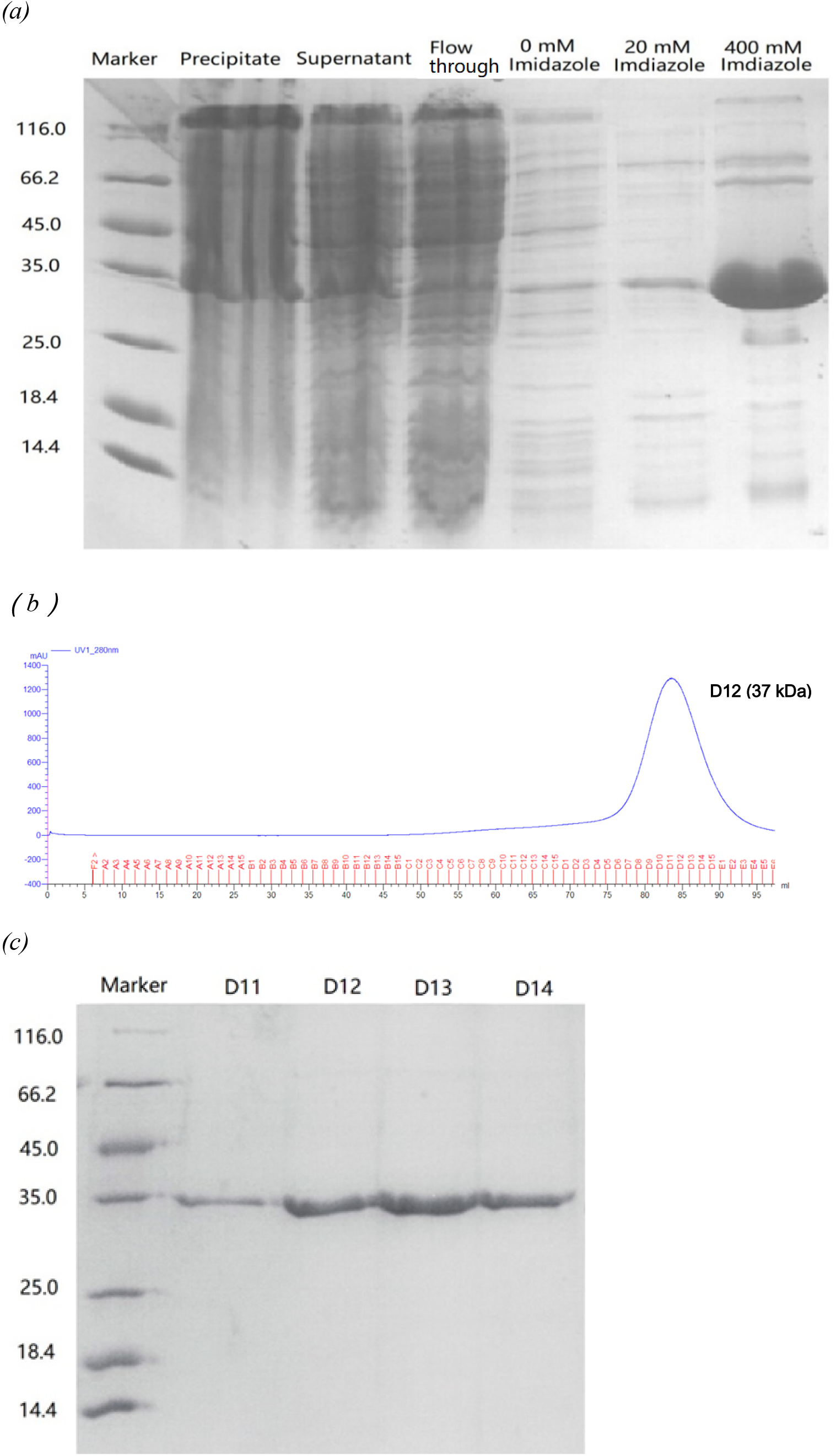
*(a)* Gel electrophoresis profile of protein fractions from centrifugation. *(b)* Gel filtration chromatography of the CbbR_7120 protein. *(c)* SDS-PAGE of the peak fractions from gel filtration chromatography.

### 3.2 Finding nick-DNAs by EMSA

The DNA sequence of rbcLXS from CbbR_7120 was estimated by the chip-seq method (Fig. 2). The FAM-tagged DNA sequence was designed based on the sequence in Table 3.

**Figure 2.**
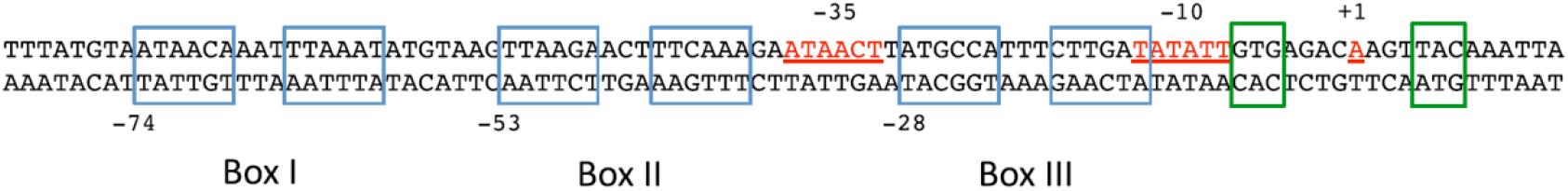
The DNA sequence of the rbcLXS promoter region^[19]^. The transcription initiation point of the operon (+1) and the −10 and −35 boxes are marked in red^[20]^. The three putative binding sites for All3953 (box I, box II and box III) are marked with blue boxes, and the NtcA-binding sites are marked with green boxes.

**Table 3.**
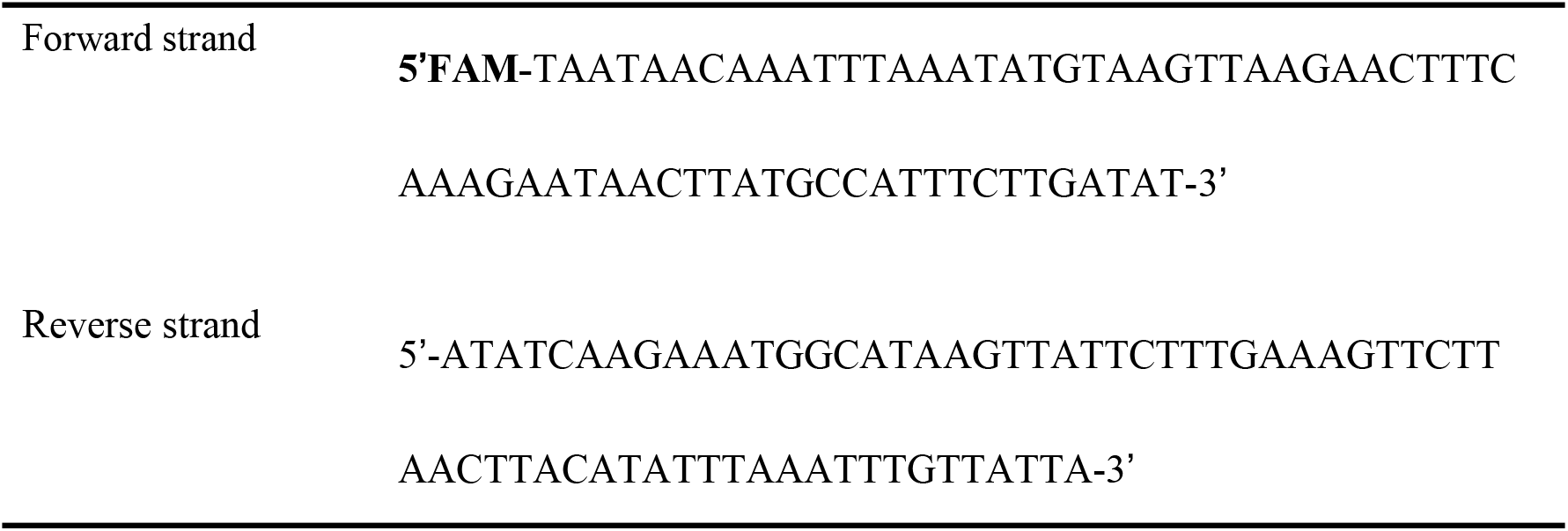
The FAM tagged DNA sequence

An amount ratio between protein and DNA equal to or greater than 5 indicates that the DNA and protein are completely bound. The concentration ratio between protein and DNA of 5 was therefore selected in the subsequent assay (Fig. 3).

**Figure 3.**
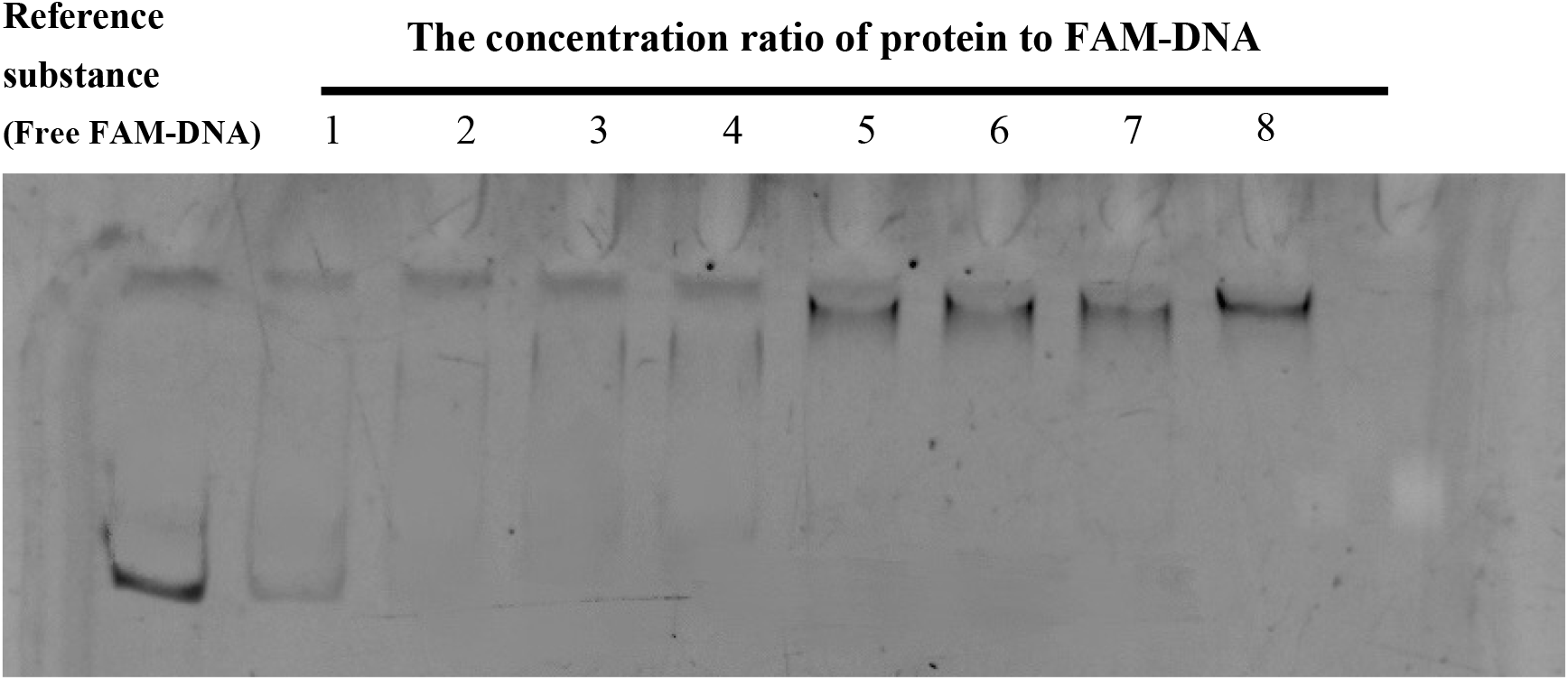
EMSA of the protein CbbR_7120 with different amounts of FAM-DNA for the minimum concentration ratio of protein-DNA-binding.

Nick-DNA fragments (BOX 1, BOX2, BOX3) that could compete with FAM-tagged DNA were designed, including BOX I, BOX II, and BOX III (Table 4); that is, three binding sites that bound to the CbbR_7120 DBD domain. This assay was designed to prove whether the speculated binding sites were correct and to confirm the binding strength of different binding sites with CbbR_7120. The protein-bound FAM-DNA was free under complete competition.

**Table 4.**
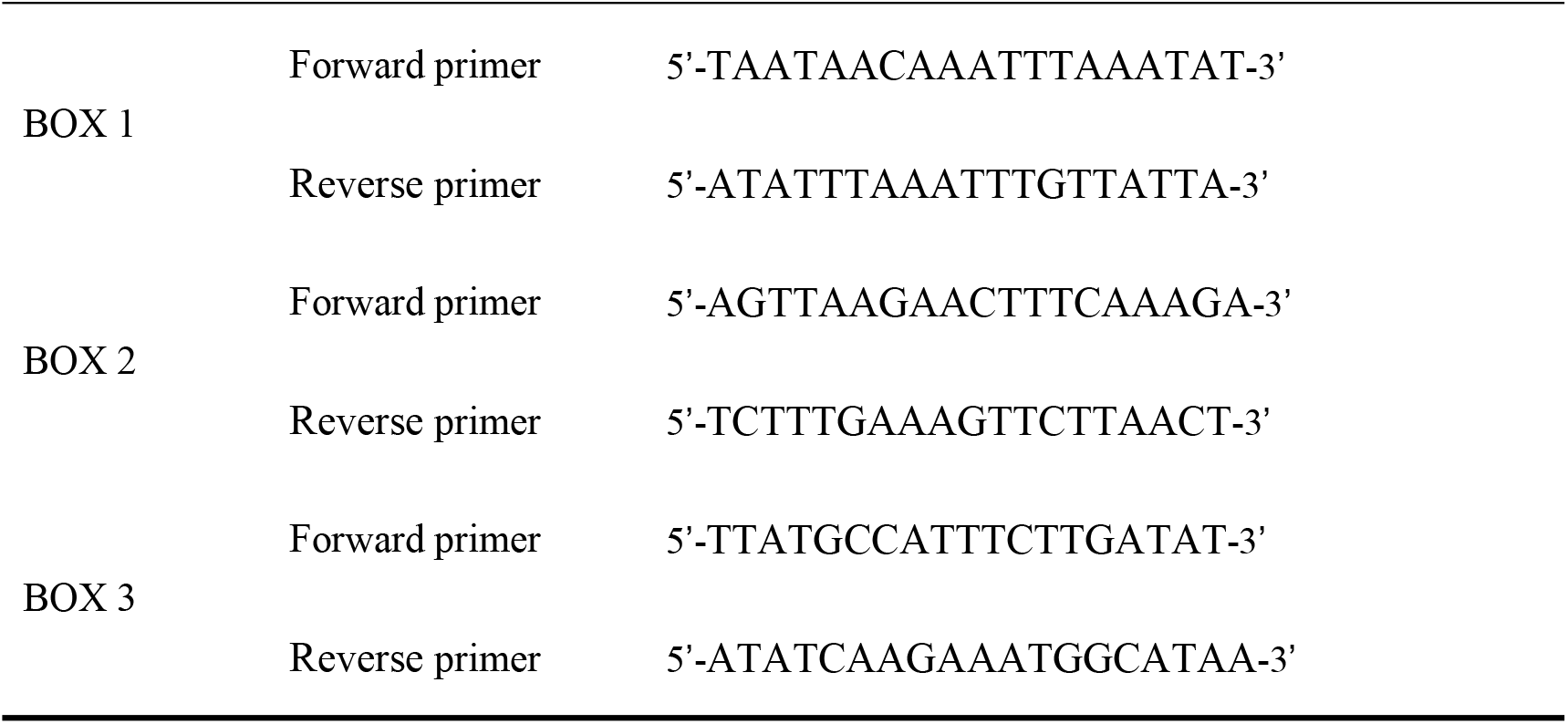
The designed competitive nick-DNA primer sequences

When competitive nick-DNA (BOX 1) competitively bound to FAM-DNA (Fig. 4a), the concentration of free FAM-DNA fragments gradually increased, while the binding band gradually weakened. The ratio between competitive nick-DNA and FAM-DNA of 400 indicated that the protein-binding FAM-DNA was free and under complete competition.

**Figure 4.**
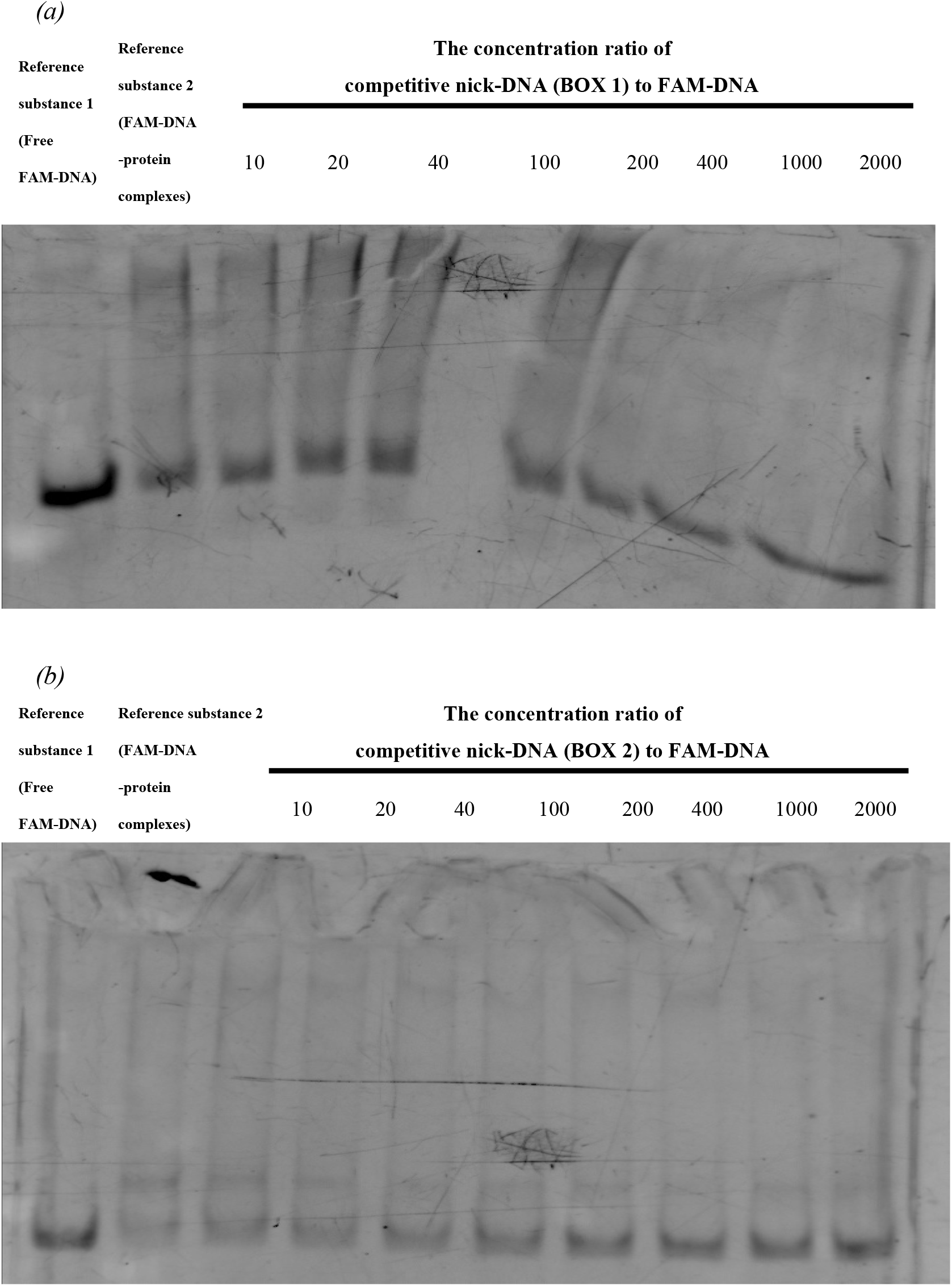
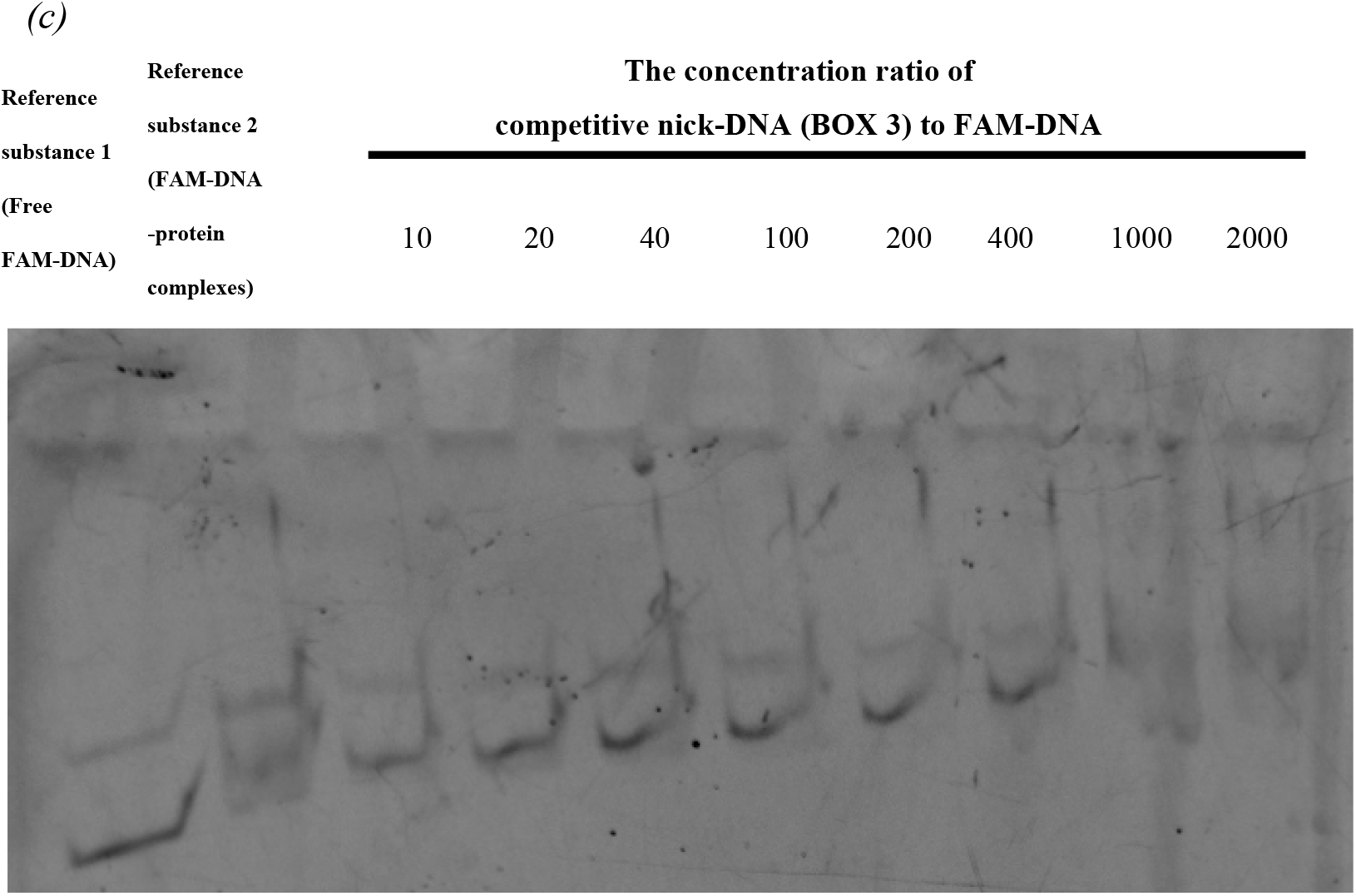
EMSA for detecting the protein binding ability of different competitive nick-DNAs with FAM-DNA.

When competitive nick-DNA (BOX 2) competitively bound to FAM-DNA (Fig. 4b), the concentration of the free FAM-DAN fragment did not significantly change due to competitive binding.

When competitive nick-DNA (BOX 3) competitively bound to FAM-DNA (Fig. 4c), the concentration of free FAM-DNA fragments gradually increased, while the binding band gradually weakened. The ratio between competitive nick-DNA and FAM-DNA of 1000 indicated that the protein-binding FAM-DNA was free and under complete competition.

Finally, it was confirmed that BOX 1 had the strongest binding ability to the CbbR_7120 protein; therefore, the seven nucleotide sequences in the linker region between BOX I and BOX II were optimized, different base-point mutations were designed from these optimal sequences, and the BOX I sequence was repeated and extended appropriately. Four optimal sequences (Box1OP1, Box1OP2, Box1OP3, Box1OP4) were formed (Table 5). Previous experiments have confirmed that the ability of BOX1OP1/BOX1OP3 to compete for binding proteins is relatively strong. Then, the concentration of competitive nick-DNA was decreased for comparison.

**Table 5.**
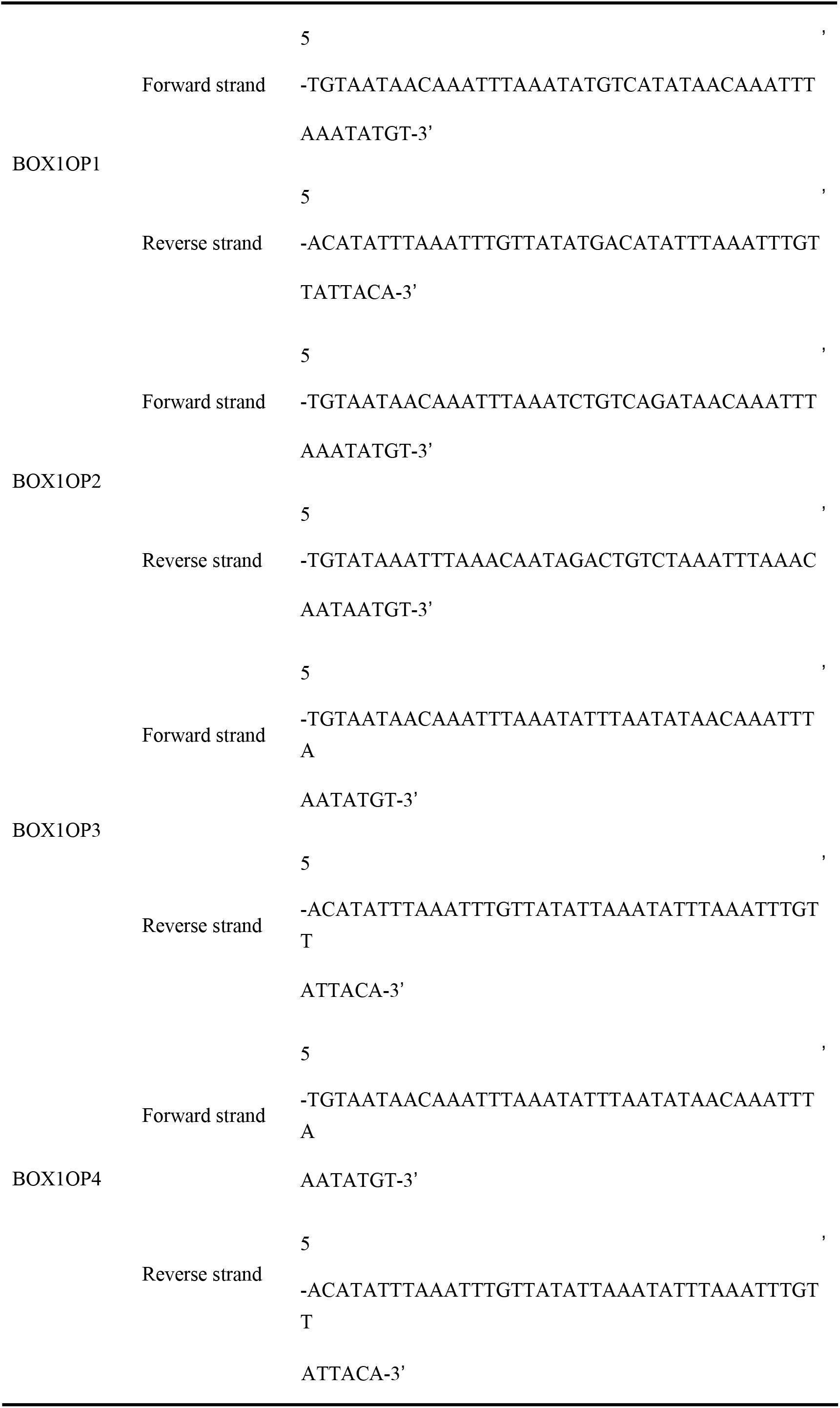
Optimized competitive nick-DNA primer sequences

When the ratio between added BOX1OP1 or BOX1OP3 and FAM-DNA concentration was 10, the binding bands of FAM-DNA with the protein generally were completely dissociated into free bands (Fig. 5). Both BOX1OP1 and BOX1OP3 had a strong binding affinity for the proteins; therefore, the two optimized fragments, BOX1OP1 and BOX1OP3, were used for the crystallization assay.

**Figure 5.**
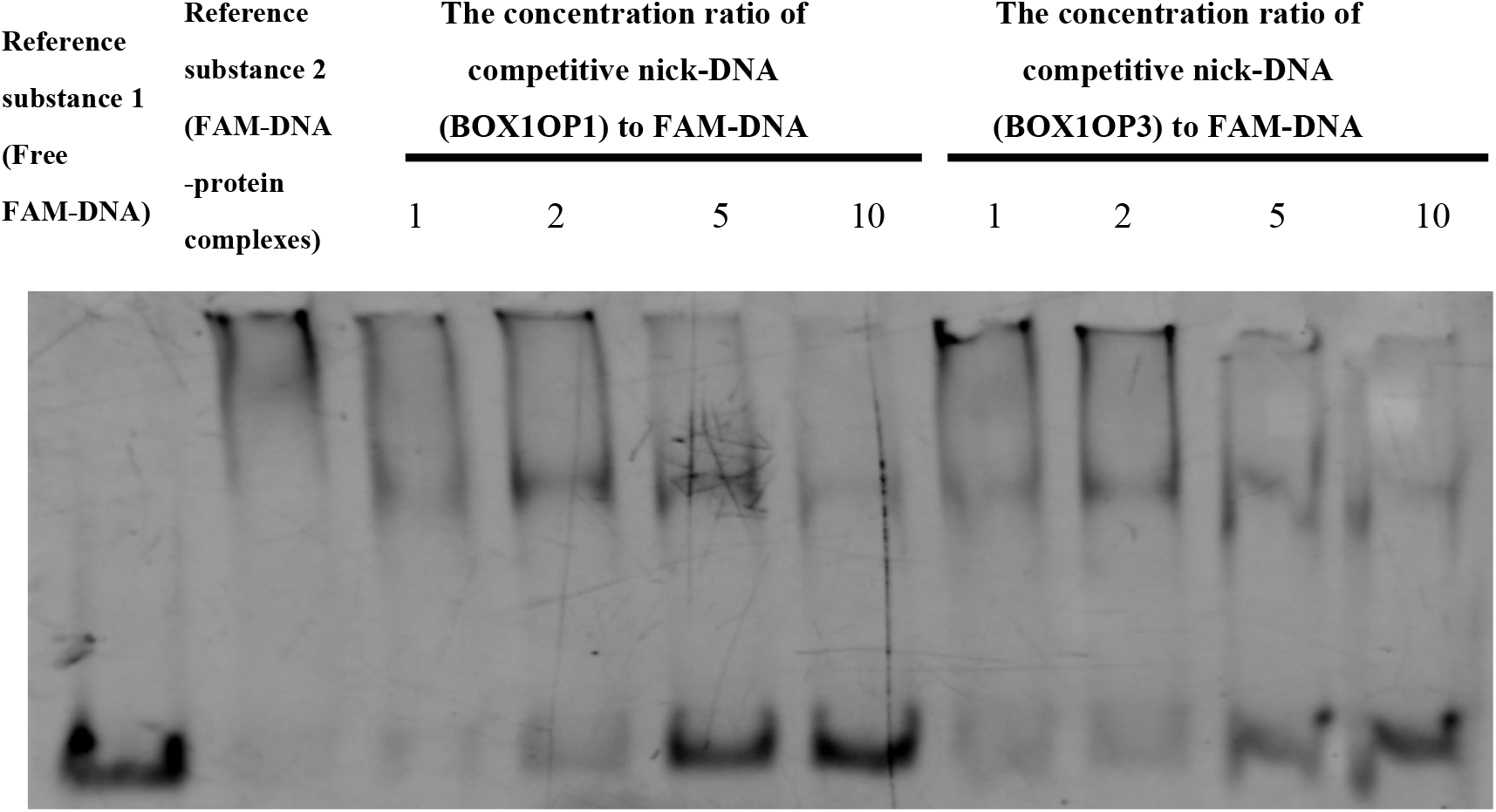
EMSA for detecting the protein binding ability of optimal competitive nick-DNAs with FAM-DNA.

### 3.3 Crystallization

CbbR_7120 was crystallized separately and then coincubated with linker DNA overnight (Table 6).

**Table 6.** Crystallization Information

| Linker_DNA | Conc. of Linker_DNA | Type of Pr. | Conc. of Pr. |
| --- | --- | --- | --- |
| BOX1OP1 | 220 mM | CbbR_7120 | 200 mM |
| BOX1OP3 | 220 mM | CbbR_7120 | 200 mM |
| - | - | CbbR_7120 | 400 mM |

Microscopic observations revealed the precipitation of protein crystals under the following crystallization conditions (Fig. 6a).

**Figure 6.**
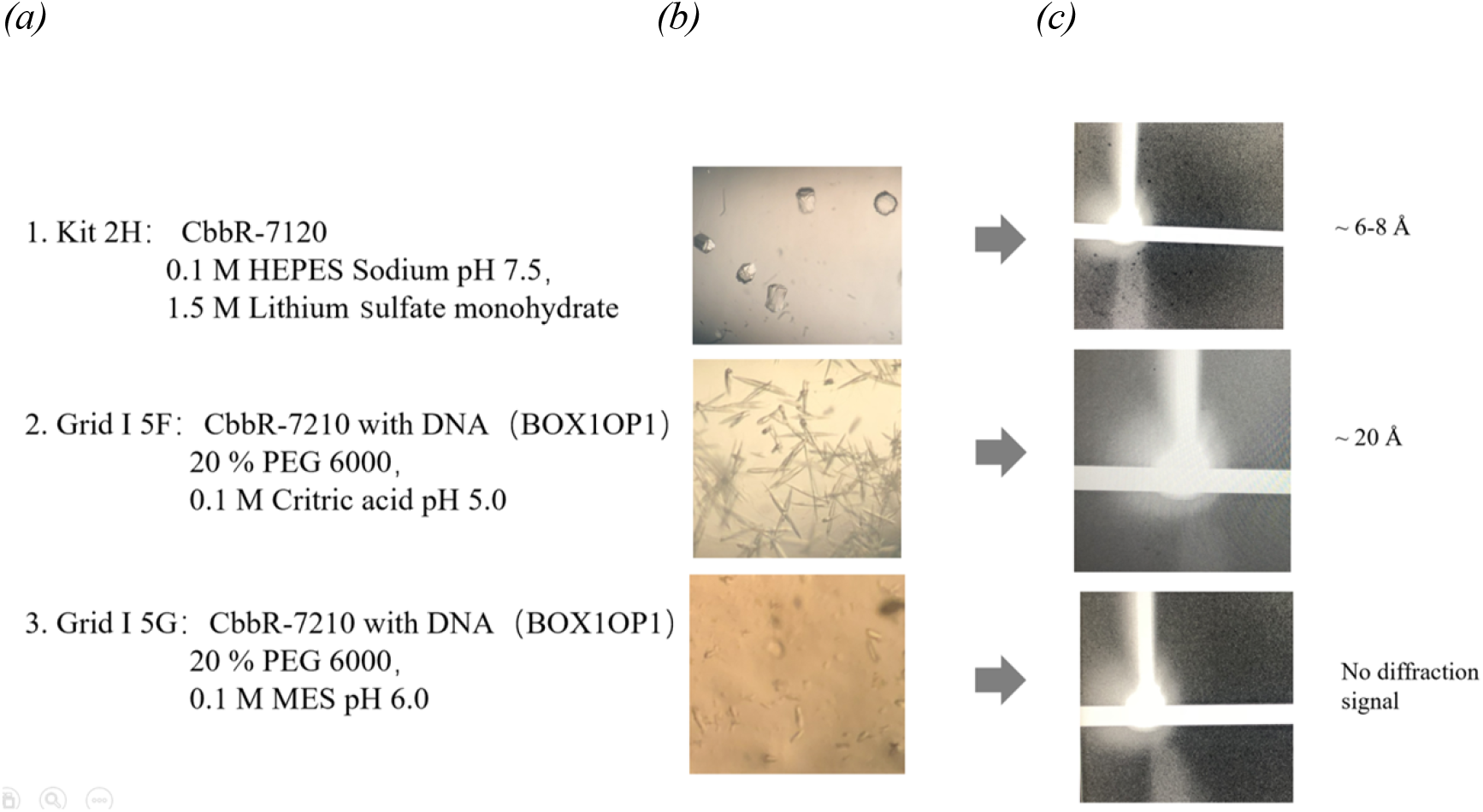
*(a)* Crystallization conditions. *(b)* Crystals of CbbR_7120 and CbbR_7120 with DNA. *(c)* X-ray diffraction patterns from the crystals.

In this study, we obtained the optimal expression conditions of CbbR in Nostoc sp. PCC 7120. The transformed, expressed and purified CbbR protein under the above conditions had high expression and relatively high purity. Additionally, an EMSA assay was used to obtain the DNA nick fragments that were tightly bound to the DBD region of CbbR_7120. Protein crystals were obtained by crystallization of the CbbR-7120, CbbR-7120 and DNA nick complexes. We optimized and analyzed the protein crystals and further elaborated their roles and functions in the CBB cycle in subsequent assays.

## Acknowledgments

This work was supported by Natural Science Foundation of Anhui Provincial Education Department (No. KJ2019A0988) and Excellent Young Talents Fund Program of Higher Education Institutions of Anhui Province (No. GXFX2017213) and Excellent Young Talents Fund Program of Higher Education Institutions of Anhui Province (No. GXYQ2019178).

